# Compensatory evolution following deleterious episodes of GC-biased gene conversion in rodents

**DOI:** 10.1101/2025.01.08.631863

**Authors:** Marie Riffis, Nathanaëlle Saclier, Nicolas Galtier

## Abstract

GC-biased gene conversion (gBGC) is a widespread evolutionary force associated with meiotic recombination that favours the accumulation of deleterious AT to GC substitutions in proteins, moving them away from their fitness optimum. In many mammals recombination hotspots have a rapid turnover, leading to episodic gBGC, with the accumulation of deleterious mutations stopping when the recombination hotspot dies. Selection is therefore expected to act to repair the damage caused by gBGC episodes through compensatory evolution. However, this process has never been studied or quantified so far. Here, we analysed the nucleotide substitution pattern in coding sequences of a highly diversified group of *Murinae* rodents. Using phylogenetic analyses of about 70,000 coding exons, we identified numerous exon-specific, lineage-specific gBGC episodes, characterised by a clustering of synonymous AT to GC substitutions and by an increasing rate of non-synonymous AT to GC substitutions, many of which are potentially deleterious. Analysing the molecular evolution of the affected exons in downstream lineages, we found evidence for pervasive compensatory evolution after deleterious gBGC episodes. Compensation appears to occur rapidly after the end of the episode, and to be driven by the standing genetic variation rather than new mutations. Our results demonstrate the impact of gBGC on the evolution of amino-acid sequences, and underline the key role of epistasis in protein adaptation. This study contributes to a growing body of literature emphasizing that adaptive mutations, which arise in response to environmental changes, are just one subset of beneficial mutations, alongside mutations resulting from oscillations around the fitness optimum.

## Introduction

GC-biased gene conversion (gBGC) is a transmission bias associated with recombination that favours G and C alleles over A and T alleles. This evolutionary force, discovered in the 2000s, results from a repair bias in meiotic recombination that leads to non-Mendelian segregation favouring G and C alleles in highly recombining regions (Eyre-Walker 1999; Galtier et al. 2001). gBGC is widespread in the living world and numerous studies have demonstrated its presence in a wide range of organisms (Duret and Galtier 2009; Pessia et al. 2012; Galtier et al. 2018) including yeasts (Mancera et al. 2008; Lesecque et al. 2013), plants (Hämälä and Tiffin 2020), hymenoptera (Wallberg et al. 2015), birds (Bolívar et al. 2016; Smeds et al. 2016; Corcoran et al. 2017; Rousselle et al. 2019) and mammals, among which primates and rodents have been the most thoroughly studied (Galtier et al. 2009; Glémin et al. 2015; Williams et al. 2015; Halldorsson et al. 2016; Li et al. 2019). gBGC has a strong influence on the evolution of patterns of genomic variation. In particular, it drives genomic base composition by increasing the frequency of G and C alleles in regions with a high recombination rate (Pessia et al. 2012; Mugal et al. 2015), as demonstrated by numerous studies in mammals (Meunier and Duret 2004; Duret and Arndt 2008; Duret and Galtier 2009; Romiguier et al. 2010; Clément and Arndt 2013; Pracana et al. 2020). A striking example is the mouse *Fxy* gene, which has experienced a spectacular increase in GC content following its recent translocation into the high-recombining pseudoautosomal region (Galtier and Duret 2007). By increasing the frequency of GC alleles and decreasing the frequency of AT alleles at neutral sites, gBGC mimics the effect of natural selection, with AT to GC mutations having a higher probability of increasing in frequency and eventually reaching fixation, as if they were positively selected. Thus, gBGC affects the site frequency spectrum (SFS), the ratio of non-synonymous over synonymous substitutions (dN/dS) and the estimation of the adaptive rate, and can confound classical tests of positive selection (Galtier and Duret 2007; Berglund et al. 2009; Galtier et al. 2009; Ratnakumar et al. 2010; Bolívar et al. 2016; Corcoran et al. 2017; Rousselle et al. 2019).

Importantly, gBGC also applies to functional sites, where it interferes with both positive and negative selection. In particular, because gBGC tends to increase the frequency of GC alleles regardless of their fitness, it promotes the fixation of deleterious AT to GC mutations that would otherwise be eliminated by purifying selection. gBGC therefore creates an additional mutation load within genomes. The existence of such a gBGC load has been studied theoretically: Duret and Galtier (2009) suggested by simulation that gBGC causes an increase from 4% to 16% of the deleterious substitution rate, while Glémin (2010) modeled how gBGC affects fitness by maintaining recessive deleterious mutations at high frequency in populations. Several studies have also empirically demonstrated the existence of a gBGC load that affects protein evolution in various organisms. Hämälä and Tiffin (2020) found that in *Arabidopsis thaliana*, protein evolution is strongly influenced by gBGC, which contributes to the accumulation of maladaptive mutations, limits the purging of deleterious mutations, and contributes to a global reduction in the efficiency of genome-scale selection. By analysing proteins with an accelerated rate of evolution, Berglund et al. (2009) and (Galtier et al. 2009) showed in primates that proteins can accumulate bursts of presumably deleterious amino-acid changes due to strong gBGC. In human populations, there is evidence for an increased load of disease-associated AT to GC mutations induced by gBGC (Necşulea et al. 2011; Lachance and Tishkoff 2014). In the mouse, after its translocation to the highly recombining pseudoautosomal region, the *Fxy* gene experienced the quick fixation of many deleterious AT to GC mutations, which would otherwise have been counter-selected (Galtier and Duret 2007).

The recombination rate varies not only among genomic regions, but also potentially in time. If the recombination map is evolutionarily unstable, such that the recombination rate of a particular locus goes up and down, we expect gBGC to be episodic. In many mammals, recombination is concentrated in hotspots, and the location of hotspots is controlled by the PRDM9 protein, which binds to specific DNA motifs (Baudat et al. 2010; Myers et al. 2010; Parvanov et al. 2010; Sandor et al. 2012). The PRDM9 gene and its binding motifs evolve very rapidly, resulting in a rapid turnover of recombination hotspots (Oliver et al. 2009; Myers et al. 2010; Berg et al. 2011; Lesecque et al. 2014; Baker et al. 2015; Latrille et al. 2017). Since gBGC is associated with recombination, this implies that gBGC should proceed in an episodic manner in mammals carrying a functional PRDM9, such as primates and rodents. Suppose that a recombination hotspot appears near a functional element (e.g. an exon). We expect a strong impact of gBGC during the episode, with an accumulation of deleterious AT to GC mutations within the concerned element, and we expect this effect to stop once the recombination hotspot dies and the gBGC episode ends. Consider a coding region affected by an episode of gBGC leading to the fixation of deleterious AT to GC mutations, thus altering the protein function. One might expect that once the recombination hotspot has died and the gene is no longer under strong gBGC, selection may act to repair the damage caused by the gBGC episode through compensatory evolution. By analysing the expected effect of gBGC on the distribution of fitness effect of new mutations, a recent theoretical study indeed suggested that gBGC episodes might considerably alter the evolutionary trajectory of the affected proteins (Joseph 2024). However, no empirical analysis of this process has been conducted so far.

Phylogenetic analyses have been carried out to identify functional sequences having undergone gBGC episodes in specific lineages (Galtier and Duret 2007; Galtier et al. 2009; Galtier 2021). This was achieved by mapping substitutions to the branches of the species tree, and identifying clusters of AT to GC changes. A similar approach might be taken to analyse the evolution of these functional sequences in downstream lineages to test whether post-gBGC episode compensatory evolution is detectable - the expectation would be an acceleration of the non-synonymous substitution rate after gBGC episodes. This requires a high enough phylogenetic resolution, such that gBGC-induced and compensatory substitutions happen in different branches of the tree and can therefore be distinguished from each other.

Murinae rodents are a particularly suitable group to study these issues, because (1) they have been shown to experience numerous and pronounced episodes of gBGC (Galtier 2021) ; (2) in rodents a large proportion of recombination hotspots are determined by the PRDM9 gene (Joseph et al. 2024) and there is a rapid turnover of these hotspots (Smagulova et al. 2016; Latrille et al. 2017) ; (3) the house mouse (*Mus musculus*) genome is particularly well known and reliably annotated; and (4) the recent sequencing of the entire exome of many Murinae species offer a large sample of proteins with a particularly high phylogenetic resolution (Roycroft, Achmadi, et al. 2021; Roycroft, MacDonald, et al. 2021).

However, it is important to note that gBGC in mammals and rodents is not always episodic. This is because the recombination rate variation is not only regulated by PRDM9, and hotspots can be determined by other means than PRDM9, even when it is present (Joseph et al. 2024). At fine scale, PRDM9-independent hotspots have been documented (Axelsson et al. 2012; Auton et al. 2013) and are often localised at the 5’ end of genes in CpG islands (Kawakami et al. 2017; Joseph et al. 2024; Raynaud et al. 2025). On a large scale, variation in recombination rate can be determined by chromosome size or telomere proximity (Stapley et al. 2017). These non-PRDM9 determinants of the localisation of recombination hotspots can create regions where gBGC can persist over time at variable spatial scales, ranging from Kb to Mb. Such a persistent effect of gBGC has been documented, for example, in the genomes of gerbils, a group close to Murinae, where the authors found large clusters of about 100 Kb of increasing GC content (Pracana et al. 2020). To study compensation, we should therefore be careful to target only gBGC events that are localised and short-lived, when it makes sense to investigate the protein response once the episode has ended. If gBGC is less precisely located and persistent in time, it is likely to be difficult to distinguish compensatory evolution from the continuing effect of gBGC.

Here, we used phylogenetic analyses of 71,040 coding exons from 48 species of Murinae rodents from the Hydromyini tribe to search for a signature of compensatory evolution after gBGC episodes have damaged coding sequences. Hydromyini is an Australasian taxon that originated about 10 Mya and is characterised by a particularly high diversification rate (Roycroft, MacDonald, et al. 2021; Roycroft, Achmadi, et al. 2021). This makes it a particularly appropriate group to carry out compensation analyses, as they provide us with a high phylogenetic and temporal resolution. By analysing synonymous substitutions, we first identified exon-specific, lineage-specific gBGC episodes. We then analysed the molecular evolution of the affected exons, as well as other exons of the same gene, after the episodes, focusing on the non-synonymous variation. We uncover ubiquitous compensatory evolution after gBGC episodes and show that protein re-adaptation in Hydromyini rodents is rapid and probably driven by the standing genetic variation.

## Results

### Overview of the data

We analysed whole-exome data from 48 species of *Hydromyini*, a highly diversified group of Australasian *Murinae*. Using the house mouse genome as a reference for orthology calling, we generated 71,040 high-quality alignments of coding exons from 15,415 genes. We only kept the exons found in at least 3 species, the average number of species per exon being 32 (Fig. 1A). The average, minimum, and maximum number of aligned exons per gene were 4.6, 1, and 67, respectively; and 12,158 genes had 3 or more aligned exons (Fig. 1B). The average exon length, GC12 and GC3, were 241 bp (exons <100pb were discarded), 49.45% and 56.11%, respectively. We reconstructed the species phylogeny (Fig. 1C) based on the concatenation of these alignments. The phylogeny was well resolved topologically, with a bootstrap support of 100% for all internal branches. A high temporal resolution was also achieved: the tree contained many short branches, with an average branch length of 0.0038 substitutions per site. The distributions of exon length, branch length, GC3 and GC12 are shown in Supplementary Figure S1A.

**Figure 1:**
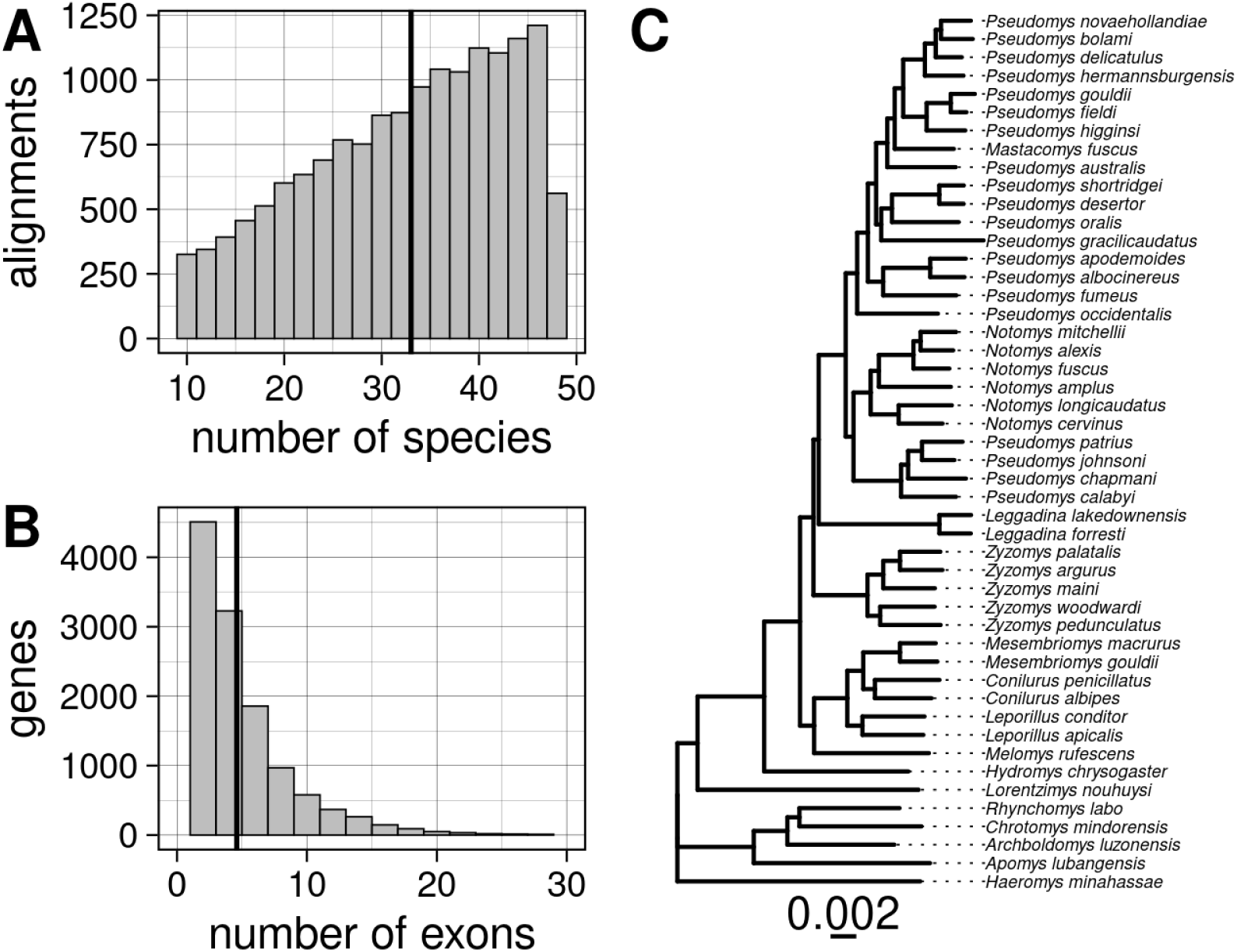
Overview of the *Hydromyini* dataset analysed in this study. **A**: distribution of the number of species per gene alignment (vertical line represents the average) ; **B**: distribution of the number of exons analysed per gene (vertical line represents the average) ; **C**: phylogeny of the 48 *Hydromyini* species, reconstructed from the concatenation of all exons.

We mapped synonymous and non-synonymous AT to GC (Weak to Strong : WS), GC to AT (SW) and GC-conservative (SSWW) nucleotide substitutions in all branches of the phylogeny, for each exon (Fig. 2A & B). We report a total of 1,855,929 synonymous substitutions and 1,320,787 non-synonymous substitutions. The counts for each type of substitution are provided in Supplementary Table S1.

**Figure 2:**
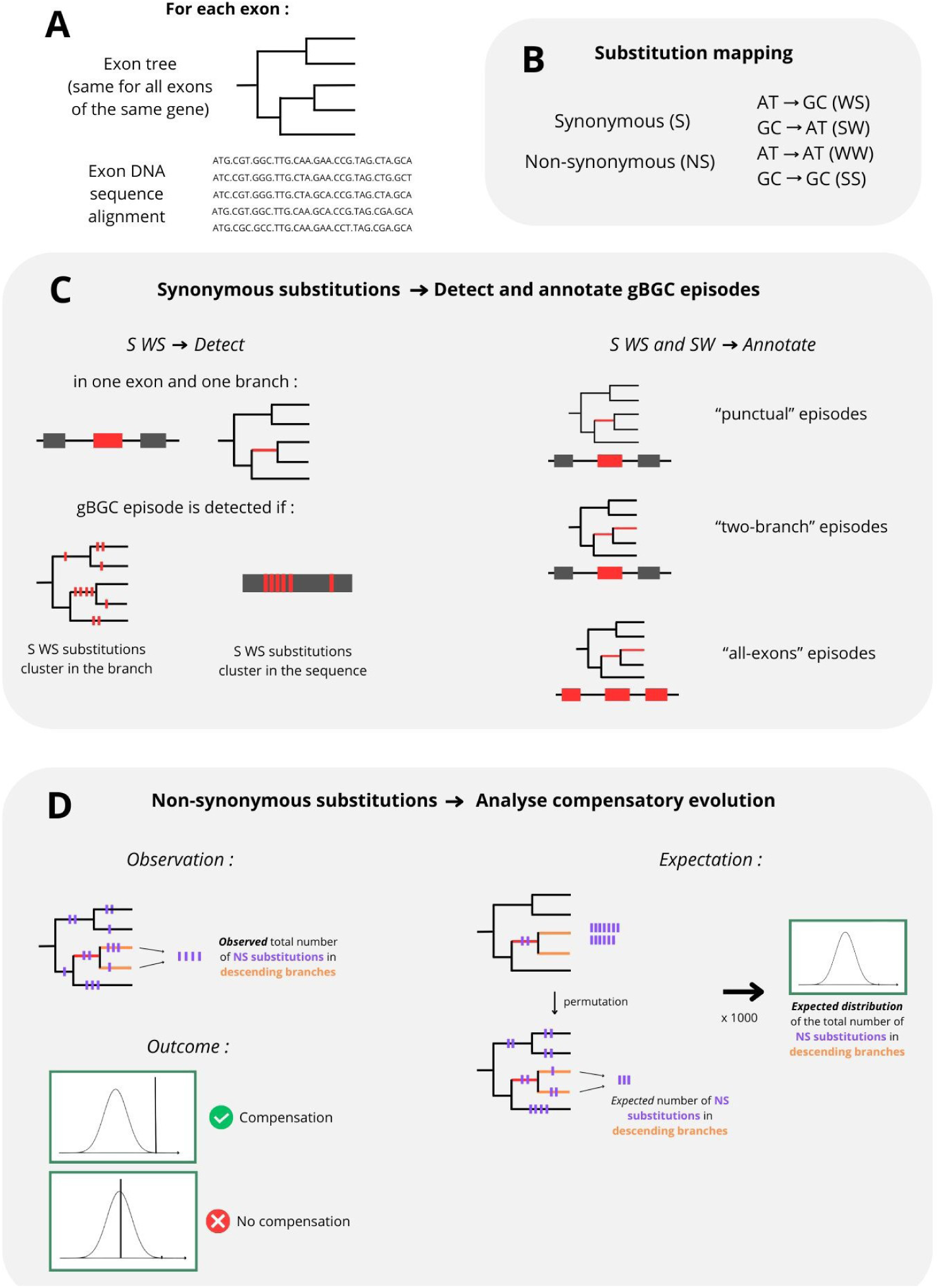
Summary of the analyses. **A**: For each exon, the data consist of a pruned tree derived from the reference phylogeny (Fig. 1C) and a corresponding DNA sequence alignment. **B**: Substitution mapping was performed for each exon, distinguishing each type of substitution. **C**: A gBGC episode (red) in a specific exon and branch was detected by identifying clusters of synonymous WS substitutions that exceed random expectations in both the tree and sequence. Based on synonymous WS and SW substitution counts across the affected branch, its descendants, the targeted exon, and other exons of the same gene, gBGC episodes were annotated as “punctual,” “two-branch,” or “all exons,” depending on their extent. Only “punctual” episodes were retained for compensatory evolution analyses. **D**: To test for compensatory evolution after gBGC episodes, the number of non-synonymous substitutions in descending branches was compared to an expected distribution generated by randomizing substitutions. A significant excess of non-synonymous substitutions indicates compensatory evolution.

### gBGC episodes

#### Detection of the episodes

Our first aim was to identify gBGC episodes by analysing the clustering of synonymous WS (S-WS) substitutions across branches and exons (Fig. 2C). Specifically, we defined a gBGC episode when we detected an excess of S-WS substitutions in a specific branch for a specific exon, compared to a random distribution of substitutions (see Material and Methods). Using this method, we detected a total of 29,845 gBGC episodes. gBGC episodes occurred in slightly longer exons and branches than the overall mean (240 bp and 0.0038 substitutions per site, respectively), with an average exon length of 267 bp and an average branch length of 0.0054 substitutions per site. The GC-content distribution of exons affected by one or several episodes was indistinguishable from the overall distribution, suggesting a homogeneous distribution of episodes across genes. The distributions of exon length, branch length, GC3 and GC12 of episodes are available in Supplementary Figure S1B. As expected, we observe an overrepresentation of S-WS substitutions compared to synonymous SW (S-SW) substitutions in most branches detected as episodes (Fig. 3).

**Figure 3:**
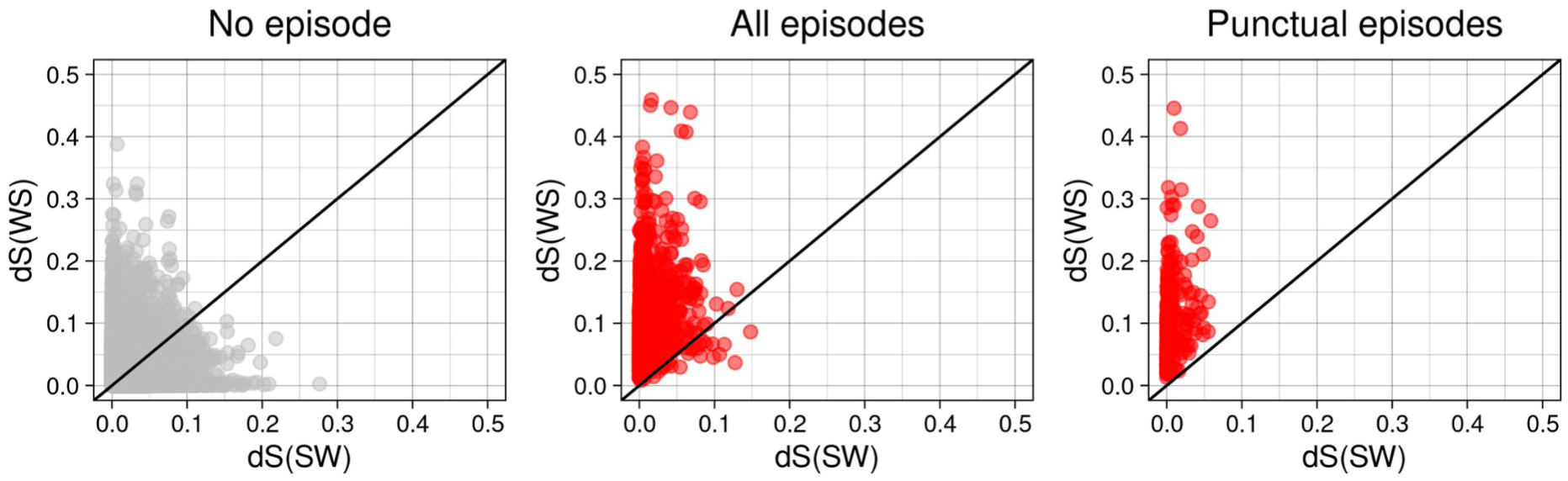
Weak-to-Strong dS (dS_WS_) and Strong-to-Weak dS (dS_SW_) in branches without (in grey) and with gBGC episodes (in red). For branches with episodes, data are shown for all types of episodes (“All episodes”) and for episodes affecting a single branch in a single exon (“Punctual episodes”), which are the subset used in compensation analyses. dS corresponds to the number of synonymous substitutions divided by the number of third codon positions, weighted by GC3 (for SW) or by AT3 (for WS). For a better visualization, we show here only a random sample of our data : 100,000 points for “No episode”, and for episodes a number of points which is proportional to the proportion of the category in all data.

#### Overflowing of the episodes and selection of punctual episodes

Our rationale involves analysing gBGC episodes that are localised and short-lived (‘PRDM9-like’ episodes), *i.e.* episodes affecting a single branch in a single exon. To check whether the detected episodes meet this criterion, we looked at the number of S-WS substitutions in the descending branches and in the other exons of the same genes. We compared the observed amount with an expected distribution obtained via 1000 randomisations of the substitutions. We observed an excess of S-WS substitutions and a deficit of S-SW substitutions in the branches immediately downstream of the episodes (Fig. 4A), as well as in the episode’s branches in the other exons of the same gene where the episodes were detected (Fig. 4B). This excess of S-WS and deficit of S-SW substitutions is a typical feature of gBGC and indicates that some of the detected episodes are not punctual, but rather extend into more than one branch and/or more than one exon.

**Figure 4:**
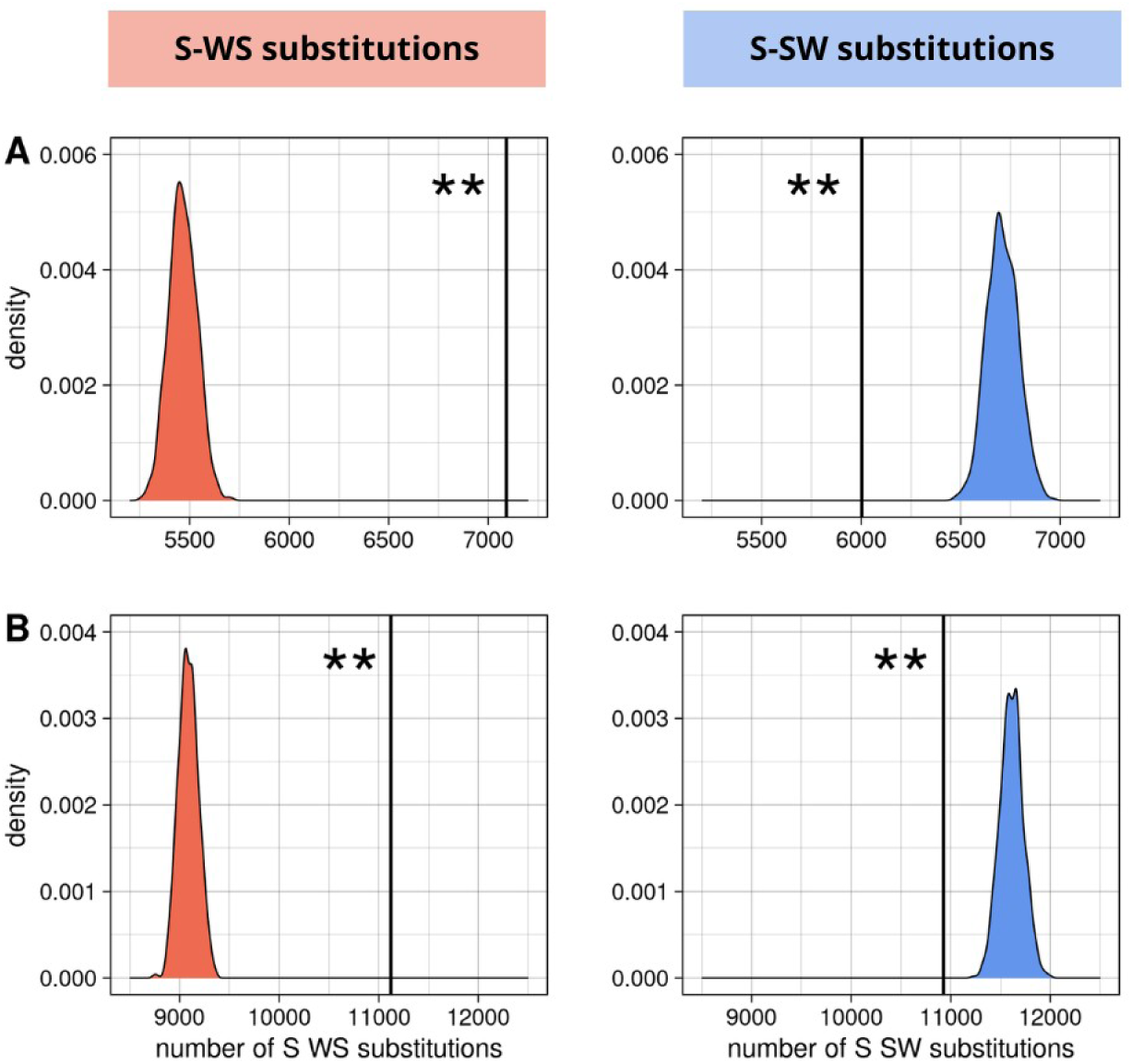
Overflowing of the gBGC episodes. **A**: in the downstream branches in the focus exon ; **B**: in the focus branches in other exons of the same gene. Vertical lines correspond to the total observed number of synonymous substitutions summed on all branches and exons concerned. Density curves correspond to the expected distribution of the total number of synonymous substitutions summed on all branches and exons concerned. ** : p-value < 0.01. Left panels (red): S-WS substitutions. Right panels (blue): S-SW substitutions.

We therefore developed an ML method (Supplementary Methods) to classify the gBGC episodes into 3 categories : punctual (localised, short-lived episodes occurring in a single exon and a single branch of the tree, the ones we are interested in), two-branch (episodes occurring in a single exon but affecting the downstream lineages) and all-exons (gene-wide gBGC) (Fig. 2C). This only concerns episodes having occurred in internal branches of the tree. 18,730 terminal-branch episodes were therefore discarded at this stage. Single-exon genes were also excluded at this stage since they do not allow distinguishing the two-branch and all-exons categories - there were 600 of them. The best-fit parameters modeled our set of detected episodes as a mixture of punctual, two-branch and all-exons episodes in proportions 0.65, 0.28 and 0.07, respectively, with an equilibrium GC3 of 94.0%, 94.6% and 84.7%, respectively. We then assigned a category to each episode according to the episode-specific posterior probabilities calculated by the method (Supplementary Methods, equations 8-10). 364 episodes (3.35%) were considered to belong to neither of these categories per their Goodness of Fit statistics (GoF < 0.2). 7,572 gBGC episodes were classified as punctual and deemed suitable for compensatory evolution analysis. Figure 3 shows that the ratio of S-SW to S-WS substitution rate tends to be higher in episodes annotated as punctual (Fig. 3C) compared to the whole set of episodes (Fig. 3B).

#### Non-synonymous evolution in “punctual” episode-carrying branches

There were 6.9 times more non synonymous WS (NS-WS) substitutions in branches carrying a punctual episode than in branches without episodes. This ratio was much lower for the other types of substitutions: it was 2.7 and 4.7 for the SW and SSWW types, respectively. The ratio remained almost unchanged after correction for exon and branch length (Supplementary Table S1). This confirms our working hypothesis that gBGC episodes are responsible for an increase in the rate of NS-WS (potentially deleterious) substitutions. 2,159 of the episodes identified as punctual were associated with at least one NS-WS substitution, while 5,413 had none.

### Compensatory evolution analyses

Our second aim was to investigate whether gBGC episodes are followed by a phase of compensatory protein evolution (Fig. 2D). To do this, we focused on non-synonymous (NS) substitutions, and analysed whether they tend to be in excess in the branches downstream of gBGC episodes. We therefore counted the number of NS substitutions in the two daughter branches of punctual episode-carrying branches, and compared it with a null distribution generated by randomising NS substitutions across exon trees. We made this comparison separately for two types of episodes : (1) those associated with at least one NS-WS substitution, *i.e.* episodes that have potentially produced deleterious substitutions after which compensation might happen; and (2) those not associated with any NS-WS substitution, *i.e.* episodes after which *a priori* no compensation is expected. Here we only considered punctual episodes occurring in an internal branch of the species tree, for which downstream lineages can be examined. There were 7,676 such episodes, of which 2,159 were associated with a NS-WS substitution. The two types of episodes did not seem to differ in terms of gBGC strength, with an average number of S-WS substitutions per site of 0.014 and 0.015 for episodes with and without NS substitutions, respectively. We carried out this analysis in the exons affected by the episodes, but also in the other exons of the same gene.

#### Signal of compensatory evolution after “punctual” gBGC episodes

In exons affected by punctual episodes, we observe a significant excess of NS substitutions after episodes with at least one NS-WS substitution, but no such excess after episodes without any NS-WS substitution (Fig. 5A). This is consistent with the hypothesis that proteins respond to deleterious gBGC episodes by a phase of compensatory evolution. Interestingly, this response is largely due to WS substitutions, and also to SSWW substitutions, but not to SW substitutions (Fig. 5B). As a control, we performed the same analyses with the two other types of episodes: the “two-branch” episodes (Supplementary Figure S2) and the “all-exons” episodes (Supplementary Figure S3), and we did not find the same signal of compensation as after the “punctual episodes”. For the two-branch episodes, we observed an excess of NS-WS substitutions in the descending branches irrespective of the presence of a substitution during the episode. This suggests that this excess of NS substitutions after “two-branch” episodes is probably not due to compensation but rather to the overflow of the episodes in the descending branches. We observed a similar signal for the “all-exons” episodes, although it is less clear, maybe due to the small sample size. We also reconducted the analysis using more stringent criteria of topological concordance (see Materials and Methods) between exon and reference trees, and found that the results were qualitatively unchanged (Supplementary Figure S4).

**Figure 5:**
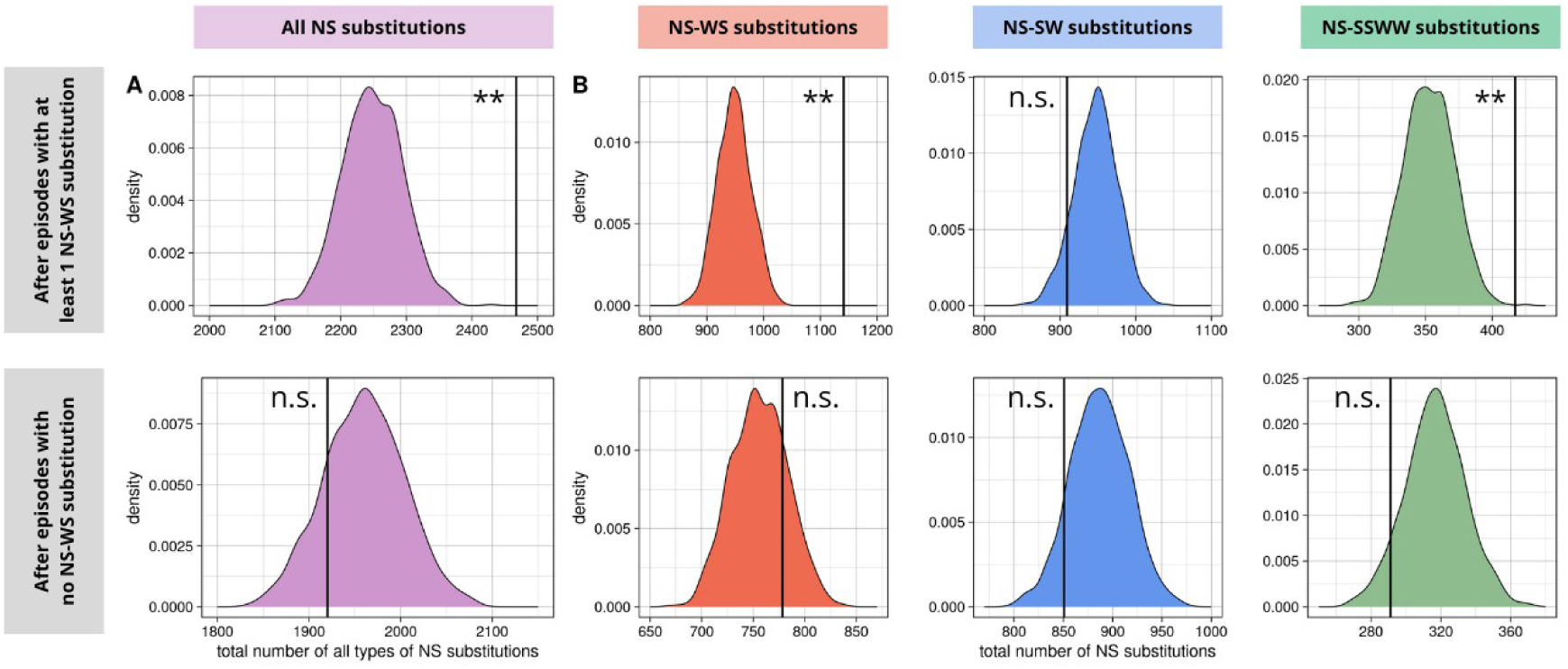
Compensation after “punctual” gBGC episodes. **A**: Total number of non synonymous substitutions observed in branches following the episodes compared with the expected distribution (summed on all exons and branches concerned). Distinction is made between episodes with at least 1 NS-WS substitution (upper panel) and episodes with no NS-WS substitution (lower panel). **B**: Same thing detailed for each of the 3 types of substitutions (red: WS, blue: SW and green: SSWW). n.s.: p-value >= 0.05 ; ** : p-value < 0.01.

#### The signal of compensation is only detectable after short branches

We reconducted the above analysis after splitting the data into 5 bins of episode-carrying branch length (Fig. 6). The results show that when episodes happen in branches longer than ∼0.006 substitutions per site, the compensation signal is no longer detectable in the descending branches. This suggests that compensation occurs quickly after the end of gBGC episodes, such that the episode and the compensatory phase can both happen in the same branch if long enough.

**Figure 6:**
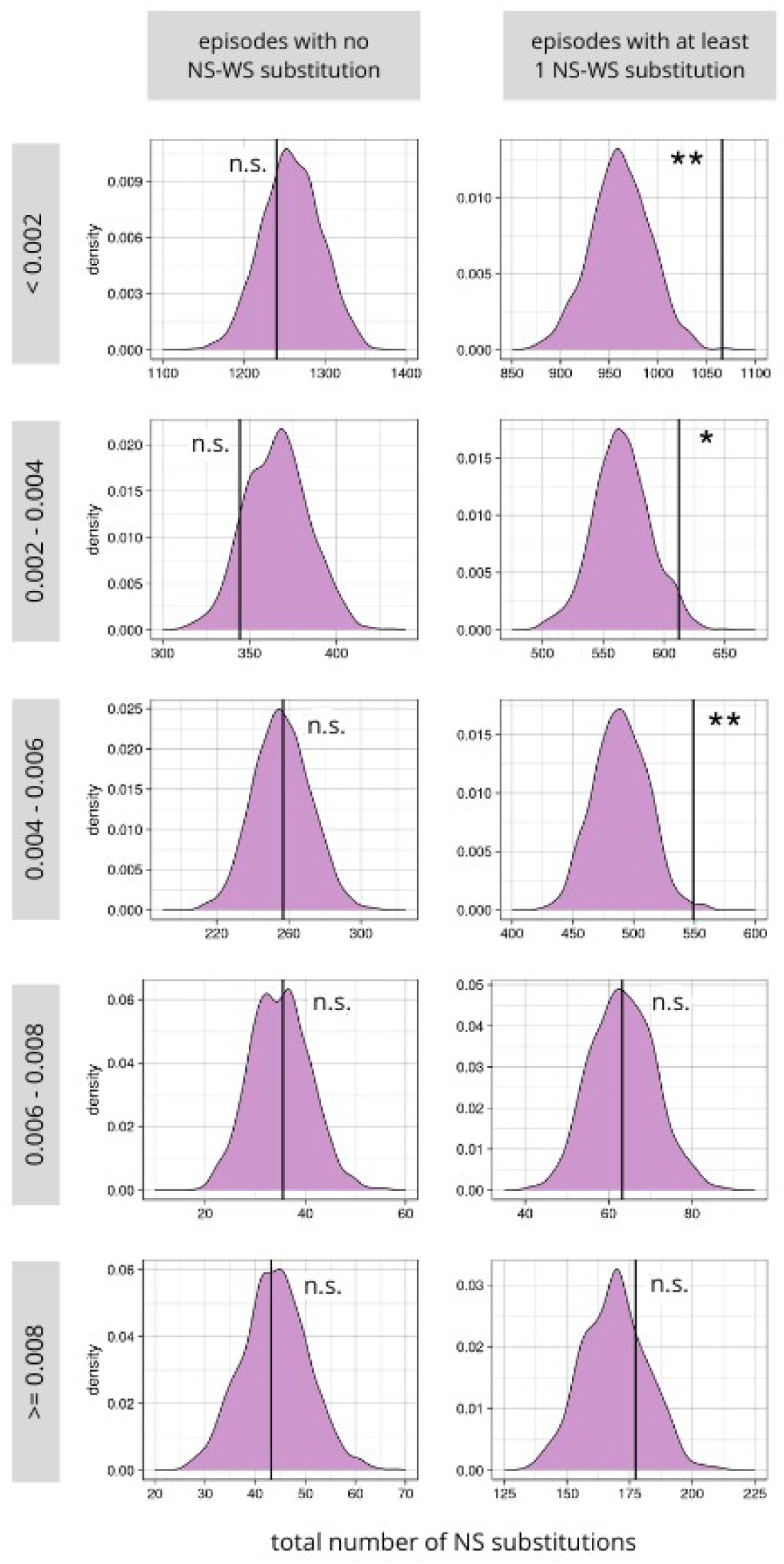
Compensation after “punctual” episodes is only detected after short carrying-episode branches. The data were separated in 5 bins of episode-carrying branch lengths: <0.002 (n=628); >=0.002 and < 0.004 (n=778), >=0.004 and <0.006 (n=434), >=0.006 and <0.008 (n=92) and >=0.008 (n=227). n.s.: p-value >= 0.05 ; *: p-value < 0.05 ; **: p-value < 0.01.

#### No detectable excess of substitutions before episodes

An excess of NS substitutions after punctual episodes with NS substitutions (Fig. 5A) is expected as soon as any process, biological or artefactual, favoring the presence (or inference) of NS substitutions in connected branches is at work. One such possibility is time-autocorrelation of substitutions, in the absence of compensatory evolution. Suppose for instance that the NS substitution rate of an exon varies in time and among clades, perhaps due to fluctuations of the overall selective pressure. In such a case, branches from clades with higher substitution rates would carry more substitutions than expected under our randomisation procedure, which might lead to excessive co-occurrence of substitutions in connected branches. To make sure the observed pattern indeed reflects a phenomenon of compensation, we conducted a similar analysis using episode ascending branches instead of descending branches. This analysis revealed no excess of NS substitutions in branches immediately upstream of punctual episodes, irrespective of the existence of a NS substitution within the episode (Supplementary Figure S5). We conclude that the excess of NS substitution in the downstream branches of an episode indeed reflects an evolutionary response to episode-associated NS substitutions, rather than a clustering of substitutions among connected branches.

#### Compensatory substitutions occur close to deleterious substitutions

We focused on predicted compensation cases where there was exactly one (supposedly deleterious) NS-WS substitution in the episode branch and exactly one (supposedly compensatory) NS substitution of any type in the descending branches (n=456 cases). We tested whether the predicted compensatory substitutions occurred at positions closer to the predicted deleterious substitution than expected by chance. This was achieved by counting the number of cases where the two substitutions were at a given distance from each other, with distances ranging from 1 to 12 codons and divided into 3 bins (Fig. 7). We found that this happened significantly more often than expected by chance for short distances (from 1 to 4 codons) (p-value < 0.01, Fig. 7), and that the signal decreased as the distance increased, suggesting that the predicted deleterious and compensatory substitutions are structurally related.

**Figure 7:**
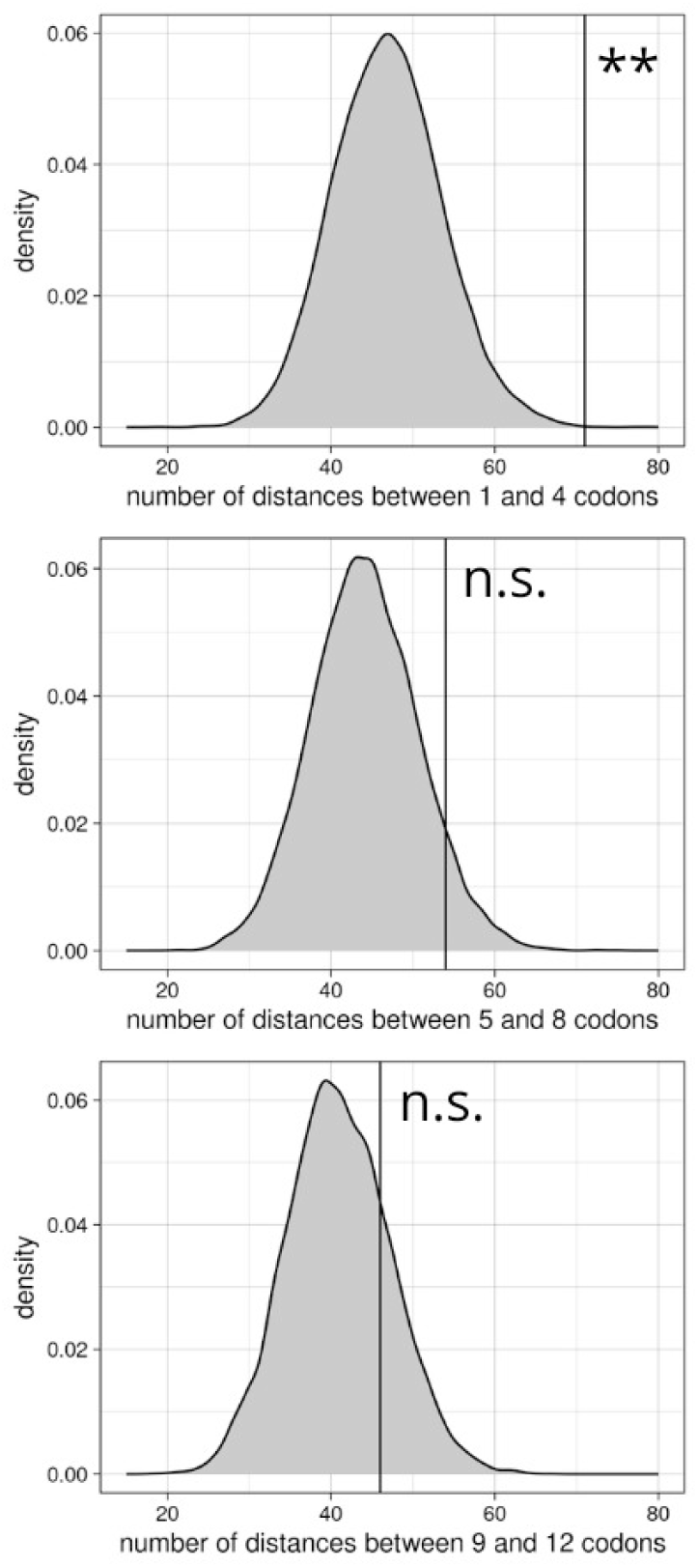
Predicted compensatory substitutions occurred at sites close to the predicted deleterious substitutions. The plots represent the total number of compensatory substitutions that are at a given distance from the deleterious substitutions, for compensatory cases where there is exactly 1 deleterious NS-WS substitution and exactly 1 compensatory NS substitution (n=456). The distances between the deleterious and the compensatory substitutions are divided into 3 bins, from upper panel to lower panel : 1-4 codons, 5-8 codons, 9-12 codons. The vertical line corresponds to the observed value and the distribution corresponds to the expected values based on 10,000 randomisations of the positions of the compensatory substitutions. **: p-value < 0.01 ; n.s.: p-value >= 0.05.

#### Compensation in the other exons during but not after punctual episodes

We also looked for potential compensation in the other exons of the same gene, during the episode (*i.e.* in the same branch) and after the episode (in the descending branches). We observed a significant signal of compensation during the episodes (Fig. 8A), but not after (Fig. 8B). We carried out the same analysis using only the other exons adjacent to the affected ones, which gave a very similar pattern.

**Figure 8:**
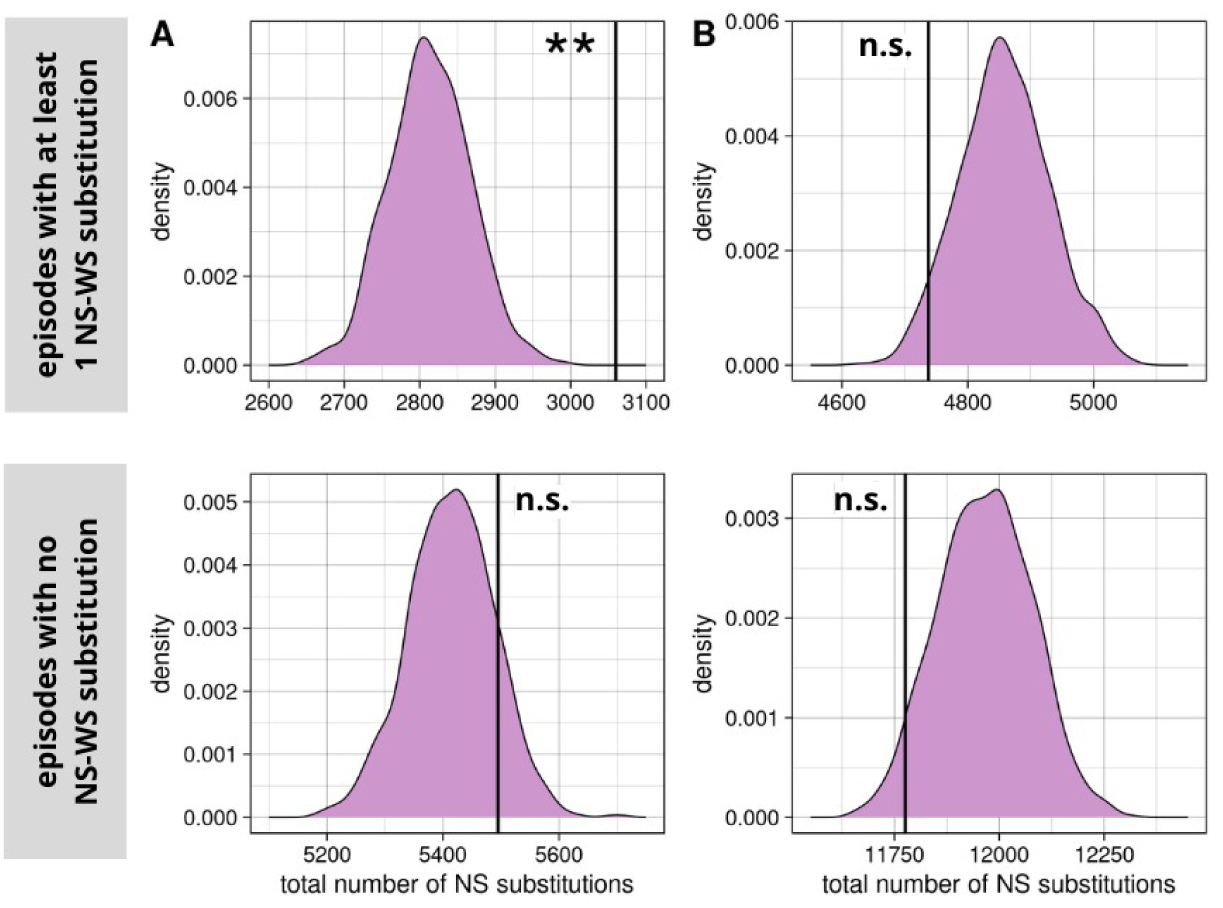
Compensation analyses in other exons of the same gene for “punctual” episodes. Observed value and expected distribution of the sum of the total number of NS substitutions of all types in other exons. **A**: during episodes (in the same branches) ; **B**: after episodes (in the descending branches). Distinction is made between episodes with at least 1 NS-WS substitution (upper panel) and episodes without NS-WS substitution (lower panel). n.s.: p-value >= 0.05 ; **: p-value < 0.01.

## Discussion

Analysing the nucleotide substitution pattern in the exon coding sequences of 48 species of Hydromyini, a highly diversified group of Murinae rodents, we detected exon-specific, lineage-specific gBGC episodes and found evidence for compensatory evolution in the descending branches.

### Prominent episodic gBGC in Hydromyini

We found that the nucleotide substitution pattern in *Hydromyini* is characterised by a clustering of WS substitutions, as found by Galtier (2021) at the Murinae scale, and by a an overrepresentation of dS_WS_ compared to dS_SW_ in most branches, as reported in the gerbil (*Gerbillinae*), brown rat (*Rattini*), and house mouse (*Murini*) lineages (Pracana et al. 2020). We identified >30,000 episodes, *i.e.*, branches with a significant excess of S-WS substitutions in a particular exon. This is much higher than the few hundreds analyzed by Berglund et al. (2009) and Galtier et al. (2009) in hominid primates. In *Hydromyini* 32% of exons and 100% of genes were affected by at least one gBGC episode. These results underscore the prominence of gBGC in this group, as previously suggested in other taxa of Murinae (Pracana et al. 2020; Galtier 2021). They confirm that *Hydromyini* are an appropriate group to study episodic gBGC and its consequences on protein evolution.

We highlight that a small but significant fraction of the detected gBGC episodes, referred to “all-exons”(about 7%), extend throughout the gene (and perhaps beyond) and persist over time - in contrast to the transient nature expected in “PRDM9-like” episodes. This suggests that in *Hydromyini,* large (>10kb) DNA segments undergo prolonged phases of GC-enrichment, as described by Pracana et al. (2020) in *Gerbillinae*. The mechanism behind this regional effect remains unclear and warrants further investigation (Brekke et al. 2023). Most of our detected episodes (around 65%) were classified as “punctual”, *i.e.*, were predicted to have occurred in a single exon and a single branch. These are suitable for studying compensatory evolution after a phase of intense gBGC. As we expected, gBGC episodes induced an increased rate of NS-WS substitutions, as previously reported in primates (Berglund et al. 2009; Galtier et al. 2009). Many of these substitutions are potentially deleterious (Glémin 2010), perhaps calling for a compensatory response.

Somewhat surprisingly, gBGC episodes were also accompanied by an increase in the rate of SW and SSWW substitutions - albeit weaker than the WS one (Supplementary table S1). We identify at least two non-exclusive explanations for this result. First, recombination is known to be mutagenic (Halldorsson et al. 2019). gBGC episodes are therefore likely associated with many new mutations of all types, which might eventually reach fixation and contribute substitutions along the same branch. Secondly, it should be recalled that genealogies, and particularly branch lengths, differ among loci due to the variance of the coalescent (Rannala et al. 2020). This is expected to create a correlation between substitution counts across categories: exon genealogies containing by chance one or several particularly long branches are likely to exhibit elevated substitution counts of all kinds in those branches.

### Post-episode compensatory evolution

Once a gBGC-induced deleterious mutation has occurred, we expect that other mutations occurring at different sites might compensate for the deleterious effect of the first mutation and eventually reach fixation, leading to an increase in the NS substitution rate after the gBGC episode.

We report a significant excess of NS substitutions after “punctual” gBGC episodes having caused a NS-WS substitutions (potentially deleterious episodes), but not after those that have not (non-deleterious episodes) (Fig. 5A). We suggest that the excess of NS substitutions after “punctual” deleterious episodes probably reflect compensatory evolution by epistasis in response to the alteration of the function of the hit protein. Compensation was detectable here thanks to the high temporal resolution of the Hydromyini tree, which contains many short branches (Fig. 1C). Indeed, the signal of compensation in the descending branches was only detectable when the branch carrying the episode was shorter than 0.006 substitutions per site (Fig. 6). Such short branches are rare in typical species trees. The rapid radiation of Hydromyini (about 50 species with a ∼10 My old common ancestor ; Roycroft, Achmadi, et al. 2021) appears to offer a particularly favorable context for the study of compensatory evolution.

We performed a number of controls to confirm that the detected excess of NS substitutions after episodes is a compensatory response to gBGC, rather than an effect of overflowing gBGC that would continue to “break up” the sequence in the descending branches. First, we note that the signal of compensatory evolution is clearly detectable only in “punctual” episodes, not in the “two-branch” and “all-exons” ones (Supplementary Figures S2 & S3). This does not suggest that gBGC overflowing explains our result, even though our assignment of episodes into the three categories is probably unperfect. Secondly, if the post-episode excess of NS substitutions were due to overflowing of gBGC, we would expect to detect it after non-deleterious episodes as well. Indeed, non-deleterious episodes do not appear weaker than deleterious ones, as they do not contain less S-WS substitutions on average, even after correcting for sequence length and GC-content. Moreover, the excess of NS substitutions was observed not only for the WS substitutions, but also for the GC-conservative SSWW substitutions, which are not expected to be directly affected by gBGC. Finally, the absence of an excess of NS substitutions in the ascending branches of deleterious episodes (Supplementary Figure S5) confirms that the signal we report in Figure 5 reflects an evolutionary response to gBGC episodes, not an exon-specific clustering of NS substitutions in specific portions of the tree due to, *e.g.*, a transient increase in the NS substitution rate.

Our analysis uses a common tree topology for all exon alignments, thus assuming that the consensus Hydromyini phylogeny (Fig. 1C) appropriately depicts the phylogeny of every analyzed exon - please note that alignments strongly departing from this assumption were filtered out, see Material and Methods. Still, it is likely that, for some alignments, the true exon phylogeny differs from the consensus tree, owing to, *e.g.*, incomplete lineage sorting or gene flow. Topological differences between the true and assumed trees are expected to affect substitution mapping, and particularly, generate erroneous inferred substitutions, possibly across connected branches. Note, however, that our analysis is unlikely to be strongly affected by the problem of false multiple substitutions at the same site - the typical consequence of exon tree/species tree discordance. Indeed, the episode-associated and compensatory substitutions we detect occur at distinct sites. It is difficult to conceive how discordance could result in mismapping a substitution to the episode-carrying branch at site X, and to a downstream branch at site Y. This does not suggest that exon tree/species tree discordance is a major confounder, as confirmed by the fact that our results were robust to an increased stringency in low-discordance exon selection (Supplementary Figure S4).

In the cases where there was exactly one NS-WS (probably deleterious) substitution in the episode branch and exactly one NS (probably compensatory) substitution in the descending branches, we showed that the downstream substitution occurred in positions close to the upstream one more often than expected by chance (Fig. 7). This is an additional argument in favour of the hypothesis of compensatory evolution in response to gBGC. Indeed, this is consistent with the report that, in a wide range of taxa covering Virus, Prokaryotes and Eukaryotes, the distance between experimentally confirmed compensatory and deleterious mutations in the protein sequence is shorter than expected by chance (Poon and Chao 2005; Davis et al. 2009). We also detected a more distant response, in other exons of the same gene (Fig. 8A). This might correspond to compensatory substitutions at sites close to the sites of the deleterious mutation in the tertiary structure of the protein (Davis et al. 2009; Ivankov et al. 2014; Storz 2018; Chaurasia and Dutheil 2022). Compensatory evolution in other exons was detected in the episode-carrying branch but not in downstream branches (Fig. 8B). Further analyses of this effect should take into account the tertiary structure of the affected proteins and the predicted effect of the supposedly deleterious and supposedly compensatory substitutions.

### New mutations or standing variation?

Positive selection on beneficial alleles for (re)adaptation can occur either from new beneficial mutations or from pre-existing (=standing) genetic variation, meaning newly beneficial alleles that were neutral or deleterious before the environmental change (in the case of adaptation), or before the change in genetic background (in our case of readaptation) (Barrett and Schluter 2008). Adaptation from standing variation is expected to be faster than adaptation from new mutations: pre-existing beneficial alleles are already present (no waiting time) and start at a higher frequency than new beneficial mutations (higher fixation probability). As a result, adaptation occurs mainly from standing variation when it occurs over short timescales (Barrett and Schluter 2008; Illingworth et al. 2012). We found that compensatory substitutions are detectable after episodes occurring in short branches, but not after episodes occurring in long branches (Fig. 6). A natural explanation of this result is that compensation happens rapidly after the episode ends (Supplementary Figure S6). According to this interpretation, when the episode-carrying branch is longer than ∼0.006, the compensatory substitutions happen in the same branch as the deleterious ones, and cannot be detected by our approach. This 0.006 threshold is roughly twice the average heterozygosity of Hydromyini exons (Supplementary Figure S1), suggesting that the response time is of the order of magnitude of the lifespan of a polymorphic mutation. Furthermore, we have shown that compensation occurs mainly via WS substitutions, and to a lesser extent SSWW ones (Fig. 5B). This is intriguing at first sight, but makes sense if one thinks of compensation from standing variation. Since a gBGC episode tends to increase the population frequency of polymorphic WS mutations, and decrease the SW ones, compensatory mutations picked up from the post-episode standing variation are likely to be of the WS kind, and unlikely to be of the SW kind. If however compensation happened mainly from new mutations, we would expect all mutation types to respond similarly. These two observations – fast response, GC-biased compensatory substitutions – argue in favor of standing genetic variation, not new mutations, being the dominating source of compensatory beneficial alleles after gBGC episodes in Hydromyini.

The relative contribution of the standing variation versus new mutations to adaptation has been studied in different ways. Experimental evolution studies, mainly in yeast, have yielded mixed conclusions, some suggesting that adaptation is mainly driven by standing variation (Burke et al. 2014), while others suggested that both sources of variation play a role (Ament-Velásquez et al. 2022). Similar conclusions about the contribution of both the standing variation and new mutations were drawn from studies that identify the origin of adaptation-associated mutations by comparing the genome sequences of a presumably adapted lineage (*e.g.* polar bear) and its presumably non-adapted source population (brown bear) (Samaniego Castruita et al. 2020; Estandía et al. 2023). Finally, a widely used method, applicable to diverse organisms, is to analyse population genomic patterns, with the assumption that adaptation from standing variation leaves a weaker pattern of selective sweep than expected from new mutations (Barrett and Schluter 2008). These soft selective sweeps have been investigated using large-scale polymorphism data analyses and found to be pervasive in various organisms, such as *D. melanogaster* (Garud et al. 2015) or humans, in which standing variation was suggested to be the dominant mode of adaptation (McManus et al. 2017; McManus et al. 2017). However, methods to distinguish between soft and hard sweeps are complex and confounded by neutral demographic processes, making such inferences tricky (Harris et al. 2018). Here we explore a new way to assess the origin of readaptation after gBGC in rodent species and find that it seems to be mainly driven by standing variation rather than new mutations, in line with previous studies having highlighted the important role of standing genetic variation in adaptation. This approach might easily be extended to other taxa, gBGC being ubiquitous, and in particular to investigate a potential effect of effective population size. If mutation limits adaptation (Rousselle et al. 2020), then small-Ne taxa such as primates might lack the genetic diversity necessary for a rapid response from standing variation, and instead resort to new mutations.

The above piece of discussion points to a process whereby a (future) compensatory mutation is already present at polymorphic stage during the episode, increases in frequency as the episode unfolds, and reaches fixation after the episode has ended. This standing variation-based scenario raises the possibility that the compensatory mutation reaches fixation not in just one of the two descending lineages, but in both. If it happened, such a convergent compensatory process would fail our detection method: because in this scenario the two descending lineages carry the same state, our method would presumably map a single substitution in the episode-carrying branch, instead of one substitution in each of the descending branches. Our approach is therefore conservative with this respect and probably underestimates the actual prevalence of compensatory evolution, to an extent that needs to be quantified.

### Epistasis vs. back mutations

Compensation is a special case of epistasis, when the fitness of an allele depends on the allelic state at other loci. The literature on epistasis is rich, with most studies on the topic agreeing that epistasis is ubiquitous and has a major impact on the long-term evolutionary trajectories of proteins (Ivankov et al. 2014; Starr and Thornton 2016; Storz 20018). There are three main ways to study epistasis in the existing literature (reviewed in Starr and Thornton (2016)). (1) Experimental studies of specific cases (few proteins) where deleterious genotypes are created and readaptation by compensatory mutations is studied in the lab (Poon et al. 2005; Filteau et al. 2015; Rojas Echenique et al. 2019; Serrano et al. 2021). (2) Search for coevolving sites across phylogenies, which works well in the case of a long-term pairwise interaction, but loses power if multiple sites are in interaction. This applies to studies identifying pairs of sites that tend to undergo substitutions in the same branches (Chaurasia and Dutheil 2022), as well as studies detecting an effect of substitutions at one site on the substitution rate at another site (e.g. Neverov et al. 2021). (3) Comparative sequence analysis, which typically showed that some disease-associated mutations in humans are fixed in other taxa, indirectly demonstrating the existence of compensatory epistasis (Kern and Kondrashov 2004; Azevedo et al. 2009). Of note these studies, while agreeing that epistasis is prevalent, are largely based on a small number of genes with well-characterised functions. Episodic gBGC, which promotes deleterious alleles throughout the genome (and also prevents their elimination by purifying selection), allowed the analysis of a large sample of proteins having undergone functional degradation. Our study thus provides a new and original perspective on the question, as we have studied at the whole exome level the readaptation of proteins temporarily displaced from their optimum. By considering gBGC-associated NS substitutions, we focus on a pool of substitutions presumably enriched in deleterious changes. This offers the opportunity of analyzing compensatory evolution irrespective of the identification of pairs of sites with correlated substitution patterns. Indeed, even a residue experiencing a single gBGC-associated deleterious substitution across the whole tree, which in turn induces a single compensatory substitution, can contribute to the signal we detect. Our results confirm that compensation by epistasis appears to be ubiquitous in protein evolution.

Experimental studies of compensation generally create highly deleterious genotypes and study protein readaptation starting far from the optimum. Poon and Chao (2005), for instance, introduced missense mutations in the φX174 bacteriophage genome, which reduced fitness by more than 50% in most mutants. In our case, the re-adapting proteins probably start relatively close to their optimal state since gBGC is a weak evolutionary force (Lartillot 2013; Galtier 2021). This enabled us to show that epistasis is pervasive even under weak selection, a situation that presumably concerns a large number of mutations, and is relevant for understanding patterns of genomic variation. This may explain why we observed a lower ratio of compensatory to deleterious mutations (not even 1:1, Fig. 5) than in experimental studies. For example, Poon and Chao (2005) reported an average of 11.8 compensatory mutations for 1 deleterious mutation.

Although our study focuses on epistatic compensation, it should be recalled that re-adaptation to the optimal state can also be achieved by back-mutations, *i.e* mutations occurring at the same site as the deleterious mutation and allowing a return to the initial state. Here we were not able to detect such back-mutations, probably missing a part of the re-adaptation. Comparing polymorphism and substitution patterns, (Latrille et al. 2024) have recently shown that back-mutations represent a substantial proportion of beneficial mutations in mammalian proteins. Moreover, a recent study has theoretically investigated the process of back-mutation after gBGC (Joseph 2024). Applying plausible amounts of gBGC to empirical fitness landscapes, this study shows that, by promoting the fixation of deleterious WS mutations and taking the protein away from its fitness optimum, gBGC significantly alters the distribution of fitness effects of future mutations, creating opportunities for beneficial back-mutations in substantial numbers. Back-mutations therefore appear to represent a significant proportion of the beneficial mutations that enable a return to the lost optimum. To quantify the relative contribution of compensatory (epistasis) vs. back-mutations in post gBGC episode protein re-adaptation is an appealing goal. If immediate, back-mutations allow the protein to return to the optimum in a single step - an efficient re-adaptation process. If however a pre-existing epistatic compensatory mutation increases in frequency or reaches fixation before the back mutation happens, then this might be less beneficial, the genetic background having changed. The dynamic of back-mutation in the context of epistasis has been theoretically investigated by McCandlish et al. (2016), who showed that the rate of back-mutation is expected to decrease with time when the site is involved in at least one epistatic interaction, and that back-mutations tend to occur either almost immediately after the deleterious substitution, or much later. The interaction between gBGC and selection might offer the opportunity to empirically test these predictions.

### Perspectives on gBGC as a confounding effect in detecting adaptation

Detection of adaptation is a widely studied topic in evolutionary biology and methods for inferring positive selection are based on the detection of rapid fixation events that result in an increased substitution rate (McDonald and Kreitman 1991). It has been largely demonstrated that gBGC can counfound these tests by promoting the fixation of deleterious WS mutations (Galtier and Duret 2007; Berglund et al. 2009; Galtier et al. 2009; Ratnakumar et al. 2010; Bolívar et al. 2016; Corcoran et al. 2017; Rousselle et al. 2019). Another previously neglected confounding effect has recently emerged in the literature, namely the confusion between beneficial and adaptive mutations (Stolyarova et al. 2020; Joseph 2024; Latrille et al. 2024). These authors point to the difference between beneficial mutations, which increase fitness, and adaptive mutations, which are the subset of beneficial mutations responding to an environmental change. In mammals, it has been quantified that non-adaptive beneficial back-mutations, which occur after a fitness decrease (*e.g.* due to drift or to gBGC), account for a large proportion (between 20% and 40%) of beneficial mutations (Latrille et al. 2024). We showed that, in Hydromyini, episodic gBGC contributes to this effect by moving proteins away from their optimum, leading to the fixation of compensatory mutations that are beneficial but not adaptive. Our study therefore demonstrates the influence of gBGC on protein evolution, and joins this early literature highlighting the non-negligible proportion of non-adaptive mutations among beneficial mutations, here focusing on compensatory mutations instead of back-mutations.

## Materials and Methods

### Data, Data cleaning, Sequence Assembly

Hydromyini whole-exome sequencing raw reads generated by Roycroft, MacDonald, et al. (2021) and Roycroft, Achmadi, et al. (2021) were downloaded from the NCBI database (references : PRJNA729818 and PRJNA705792). This dataset is composed of 48 species, of which 5 are extinct. Reads came from a single individual in 42 species, from 2 individuals in 3 species, and from 3 individuals in 3 species. Raw reads were quality trimmed using Trimmomatic version 0.39 (Bolger et al. 2014) and pooled for the species with several individuals. Cleaned reads were assembled using TRINITY version 2.13.2 (Grabherr et al. 2011) to obtain a set of contigs of at least 200 bp for each species.

### Orthologous exon call

To identify orthologous coding exon sequences in our assembled contigs, we downloaded house mouse reference coding exons from the Ensembl database (‘Ensembl genes 109’ - ‘Mouse genes GRCm39’) and used a reciprocal best BLAST-hit strategy. First, the reference exon sequences were blasted against our contig sequences with the blastn function of the BLAST tool version 2.12.0 (Altschul et al. 1990), requiring >85% sequence identity, an alignment length >=100bp and covering at least 50% of the reference exon length. The first hit contigs were trimmed by only keeping the parts included in the BLAST alignments. Finally, a reciprocal blast of the trimmed contigs against the reference exon set was performed, and we only kept the contigs hitting the same exon that hit them in the first place.

### Alignment and alignment filtering

For each reference exon, orthologous sequences were aligned with the OMM-MACSE pipeline version 12.01 (Scornavacca et al. 2019), which produces a codon-aware alignment of nucleotides using MACSE version 2.08 (Ranwez et al. 2018) and applies HmmCleaner version 1.8 for filtering out dubious parts of the alignment (Di Franco et al. 2019). Alignments were further filtered in order to exclude suspicious sequences. We mapped raw reads to our exons with bwa-mem version 0.7.17-r1188 (Li 2013) and calculated the per sequence average coverage depth and heterozygosity using samtools version 1.19.2 (Li et al. 2009) and Angsd version 0.930 (Korneliussen et al. 2014), respectively. Sequences with an average coverage depth below 5X or greater than the species mean plus 4 standard deviations were excluded. Sequences with an heterozygosity greater than the species mean heterozygosity plus 3 standard deviations were excluded, as this may be indicative of hidden paralogy (Roycroft, Achmadi, et al. 2021). We also excluded from the alignments the sequences that contained stop codons. After these filtering steps, a total of 71,040 exon alignments from 15,415 genes were retained.

### Phylogeny

A species tree was reconstructed with IQ-TREE version 2.0.7 (Minh et al. 2020) from the concatenation of all filtered alignments, using a GTR model and 1000 ultrafast bootstrap replicates. The tree was rooted with the Ape R package, version 5.8 (Paradis and Schliep 2019) using *Haeromys minahassae* as the outgroup, in agreement with Roycroft, Achmadi, et al. (2021).

We also created one tree per gene, because we need for our analyses that all exons of a same gene have the same topology. To do that, for each gene *g*, we first identified the subset of species for which a sequence of each exon of *g* was available, then we obtained the gene tree by pruning the consensus phylogeny, only keeping species from this subset (Ape “prune_tree” function).

### Topological discordance filtering

Our analysis assumes that the consensus tree appropriately represents the phylogeny of the analyzed exons. This might be untrue for some of the exons. In particular, exon alignments containing paralogous sequences are likely to depart from this assumption. To filter them out, for each exon alignment, we first reconstructed a *de novo* tree using IQ-TREE, and calculated the sum of its branch lengths. Then, still using IQ-TREE, we forced the consensus topology, re-estimated branch lengths, and again calculated their sum. We discarded exon alignments for which the constrained total tree length (consensus topology) was 1.5 times as long, or more, as the unconstrained one (free topology). Here we reasoned that an alignment strongly rejecting the consensus topology should manifest itself by a large number of homoplasic substitutions when the consensus tree is forced. Other thresholds were tried for this filter, which did not alter the results (Supplementary Figure S4).

### Substitution mapping

We realised a substitution mapping for each exon alignment using the BppSuite (Guéguen et al. 2013). We used the BppML program version 3.0.0 to optimise gene tree branch lengths under the YN98 codon model (Yang and Nielsen 1998). Then, we estimated the number of synonymous and non-synonymous substitutions that have occurred in each branch at each codon for each alignment for the four different mutational categories (WS, SW, SS, WW) in an empirical Bayesian framework using the MapNH program version 2 (Romiguier et al. 2012) with the parameter “map.type = Combination(reg1=DnDs, reg2=SW)”.

### Detection of gBGC episodes

A signature of episodic gBGC is an aggregation of WS substitutions in time and space. We used these two criteria to define and detect gBGC episodes in specific branches and specific exons. We calculated for each exon in each branch of the tree the total number of S-WS substitutions, and performed permutations to determine if this number is greater than expected in the absence of gBGC. A permutation was achieved by redistributing the total number of S-WS substitutions of an exon across the gene tree, the probability for a substitution to be assigned to a particular branch being proportional to the length of that branch. This was performed 1000 times to give, for each exon, the expected distribution of the number of S-WS substitutions in every branch of the tree in the absence of gBGC. An episode of gBGC was called in a given exon in a given branch when the observed number of S-WS substitutions in the branch was in the top 1% of the expected distribution.

For exons longer than 1000 bp, we added a criterion of spatial aggregation. We calculated Moran’s I, a metric already used by Galtier (2021) to detect gBGC, to measure the spatial aggregation of S-WS substitutions across the sequence length. We generated 1000 permutations by redistributing the substitutions occurring in a specific branch along the exon sequence. This provided us with the expected distribution of Moran’s I expected under the hypothesis of a random spatial distribution of S-WS substitutions. An episode of gBGC was called when the observed Moran’s I was in the top 1% of the expected distribution.

### Selection of ‘PRDM9-like’ episodes

We developed a maximum likelihood (ML) classifier of gBGC episodes in order to distinguish localised, short-lived episodes from large-scale and/or long-lasting effects of gBGC. Briefly, we model our set of episodes as a mixture of three categories: punctual (episodes only hitting one exon in one branch of the tree), two-branch (episodes hitting a single exon but affecting downstream lineages) and all-exons (gene-wide gBGC). The model parameters were the proportion of the three categories and the WS/SW balance at synonymous positions in the branch carrying the episode and its two descending branches. Episodes occurring in a terminal branch were therefore disregarded. Having fitted the model by maximising the likelihood, the posterior probability for a given episode to belong to a particular category was calculated in the empirical Bayesian framework. An episode-specific measure of the goodness of fit was also calculated by comparing the likelihood of this and a degenerate model. The details of the method are presented in the Supplementary Methods. We selected episodes for which the posterior probability of the punctual category was higher than the other two categories, and the goodness-of-fit *p*-value was above 0.2.

### Analysis of compensatory evolution

Having called gBGC episodes based on synonymous substitutions, we focused on non-synonymous substitutions in downstream branches to analyse the protein response. We reasoned that, if there is compensatory evolution, one should observe a surplus of NS substitutions in the two branches immediately downstream of the branch carrying the episode. Permutations were performed 1000 times for each exon to redistribute the NS substitutions of each type (WS, SW, SS, WW) across branches, blocking branches where gBGC episodes were detected, *i.e.*, the NS substitutions in these branches were not redistributed, and the redistributed substitutions could not be assigned to these branches. Observed values and expected distributions were summed across all branches immediately downstream of episodes in all exons touched by an episode. This allowed us to compare the observed number of NS substitutions in branches after episodes with its expected distribution in the absence of compensatory evolution, controlling that no episode is detected in these branches. We distinguished two types of episodes : (1) those having produced at least one NS-WS substitution, potentially deleterious, after which compensatory evolution might happen, and (2) those with no NS-WS substitution, *i.e* after which there is *a priori* no compensation to be expected. We searched for such compensation in the exons which have endured the gBGC episodes and also in the other exons of the same gene. We also looked if there is compensation during the episode in the other exons.

We then focused on the cases of potential compensation where there is exactly 1 (supposedly deleterious) NS-WS substitution in the episode branch and exactly 1 (supposedly compensatory) NS substitution of any type in the descending branches. We looked at the distance (in number of codons) between these two substitutions and assessed whether the compensatory substitutions are closer to the deleterious substitutions than would be expected by chance. To do this, the positions of the compensatory substitutions were randomised 1000 times, by drawing an integer between 1 and the number of codons of the considered exon. We then counted the number of cases where the observed distance between the deleterious and the compensatory substitutions were less than or equal to 3 codons, and compared it with the expected distribution of the same counts obtained from the randomised positions.

## Data accessibility

All the scripts and analyzed data files are available from https://github.com/Marie-98/gBGC_comp *and 10.5281/zenodo.14534842*

## Acknowledgments

We wish to thank Khalid Belkhir, Benjamin Penaud, Vincent Ranwez and Iago Bonnici for their advices and help with bioinformatics ; Laurent Duret, Marie Sémon, Thibault Latrille, Julien Joseph, Stéphanie Bedhomme and two anonymous reviewers for insightful feedback and discussions ; and Emily Roycroft for her advice on data filtering.

