## Supplementary_material for "Compensatory evolution following deleterious episodes of GC-biased gene conversion in rodents"

### Supplementary material to « Molecular re-adaptation : compensatory evolution following deleterious episodes of GC-biased gene conversion in rodents »

**Table S1 : Summary of the substitutions mapping.** The ratios of the mean number of substitutions between branches with episodes and branches without episodes, and between branches after all types of episodes and branches after punctual episodes, do not change substantially after correction for sequence length, branch length and GC content.

| | total number in all data | mean number in branches with punctual episodes | mean number in branches without episodes | mean number in branches after episodes (all types) | mean number in branches after punctual episodes |
|---|---|---|---|---|---|
| **S WS** | 759,754 | 2,94 | 0,11 | 0.42 | 0.25 |
| **S SW** | 961,523 | 0,31 | 0,14 | 0.29 | 0.28 |
| **S SSWW** | 134,652 | 0,12 | 0,021 | 0.059 | 0.044 |
| **NS WS** | 489,249 | 0,54 | 0.080 | 0.19 | 0.14 |
| **NS SW** | 624,195 | 0,16 | 0.092 | 0.15 | 0.12 |
| **NS SSWW** | 207,343 | 0,11 | 0.032 | 0.066 | 0.050 |

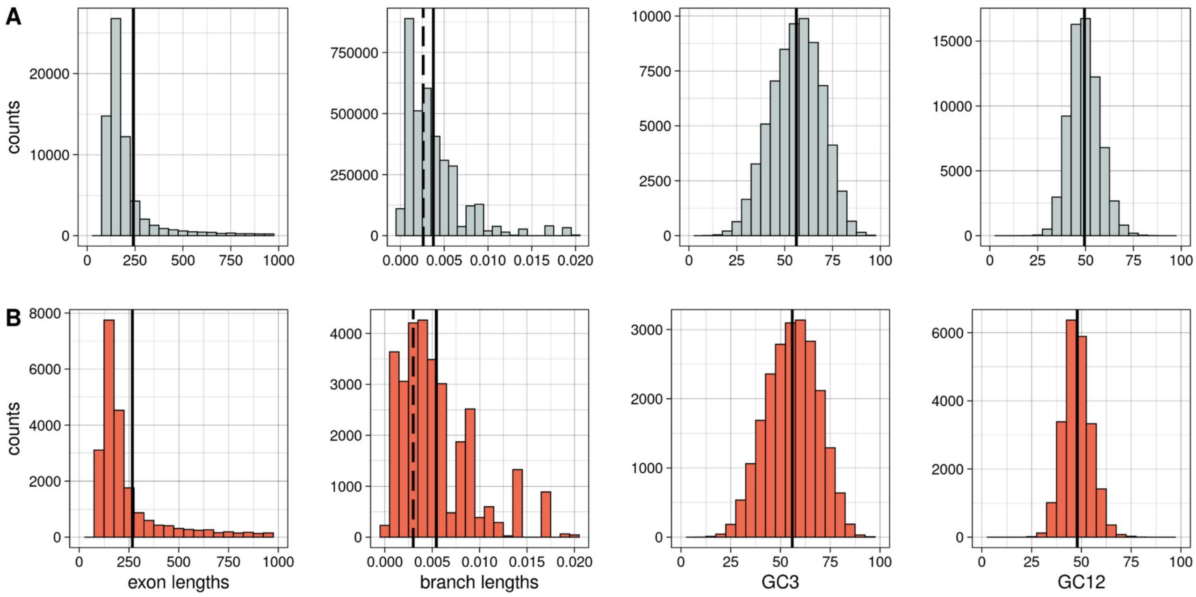


**Figure S1 : Distributions of exon length, branch length, GC3 and GC12. A** : all the data and **B** : gBGC episodes. Vertical solid lines represent the means and vertical dashed lines on the distribution of branch lengths represent the average heterozygosity.

**Table S2 : Parameter estimates of the episode's classifier model.**

| | |
|---|---|
| $p_1$ | 64.9% |
| $p_2$ | 28.0% |
| $p_3$ | 7.1% |
| $\theta_1$ | 49.5% |
| $\theta_1$* | 94.0% |
| $\theta_2$ | 68.3% |
| $\theta_2$* | 94.6% |
| $\theta_3$ | 84.7% |
| maximum ln(L) | -49924.129438 |

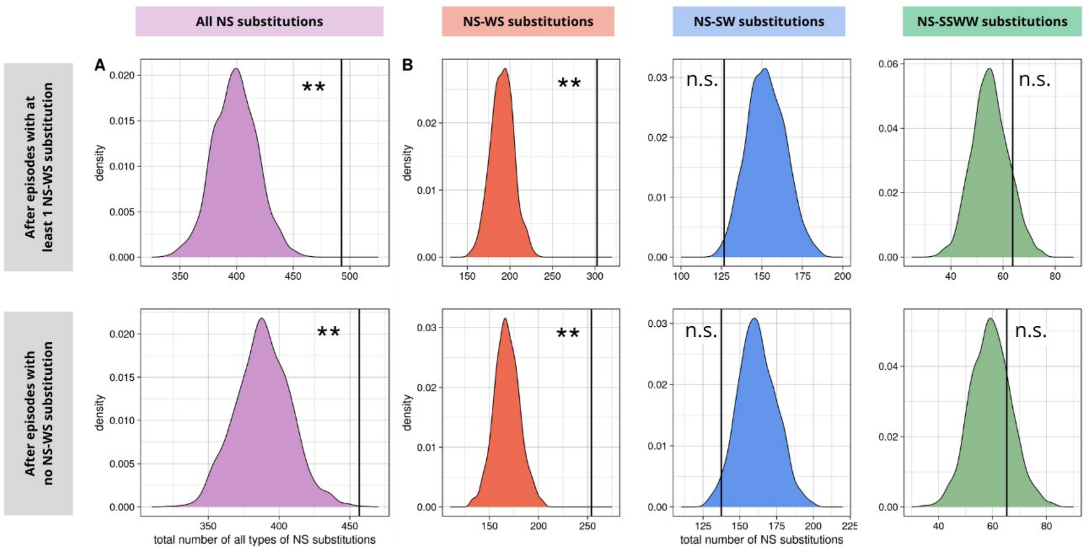


**Figure S2 : No compensation after "two-branch" gBGC episodes. A**: Total number of non-synonymous substitutions observed in branches following the episodes compared with the expected distribution (summed on all exons and branches concerned). The expected distributions are based on 1000 randomisations. Distinction is made between episodes with at least 1 NS-WS substitution (up) and episodes with no NS-WS substitution (down). **B**: Same thing detailed for each of the 3 types of substitutions (red: WS, blue: SW and green: SSWW). n.s. : p-value >= 0.05 ; ** : p-value < 0.01.

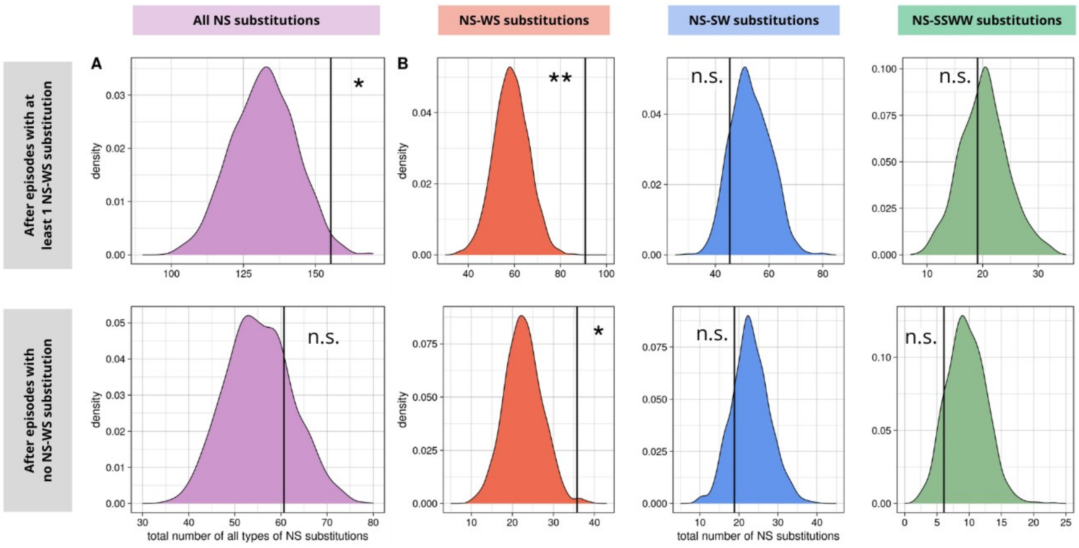


**Figure S3 : No compensation after "all-exons" gBGC episodes. A**: Total number of non-synonymous substitutions observed in branches following the episodes compared with the expected distribution (summed on all exons and branches concerned). The expected distributions are based on 1000 randomisations. Distinction is made between episodes with at least 1 NS-WS substitution (up) and episodes with no NS-WS substitution (down). **B**: Same thing detailed for each of the 3 types of substitutions (red: WS, blue: SW and green: SSWW). n.s. : p-value >= 0.05 ; * : p-value < 0.05 ; **: p-value < 0.01.

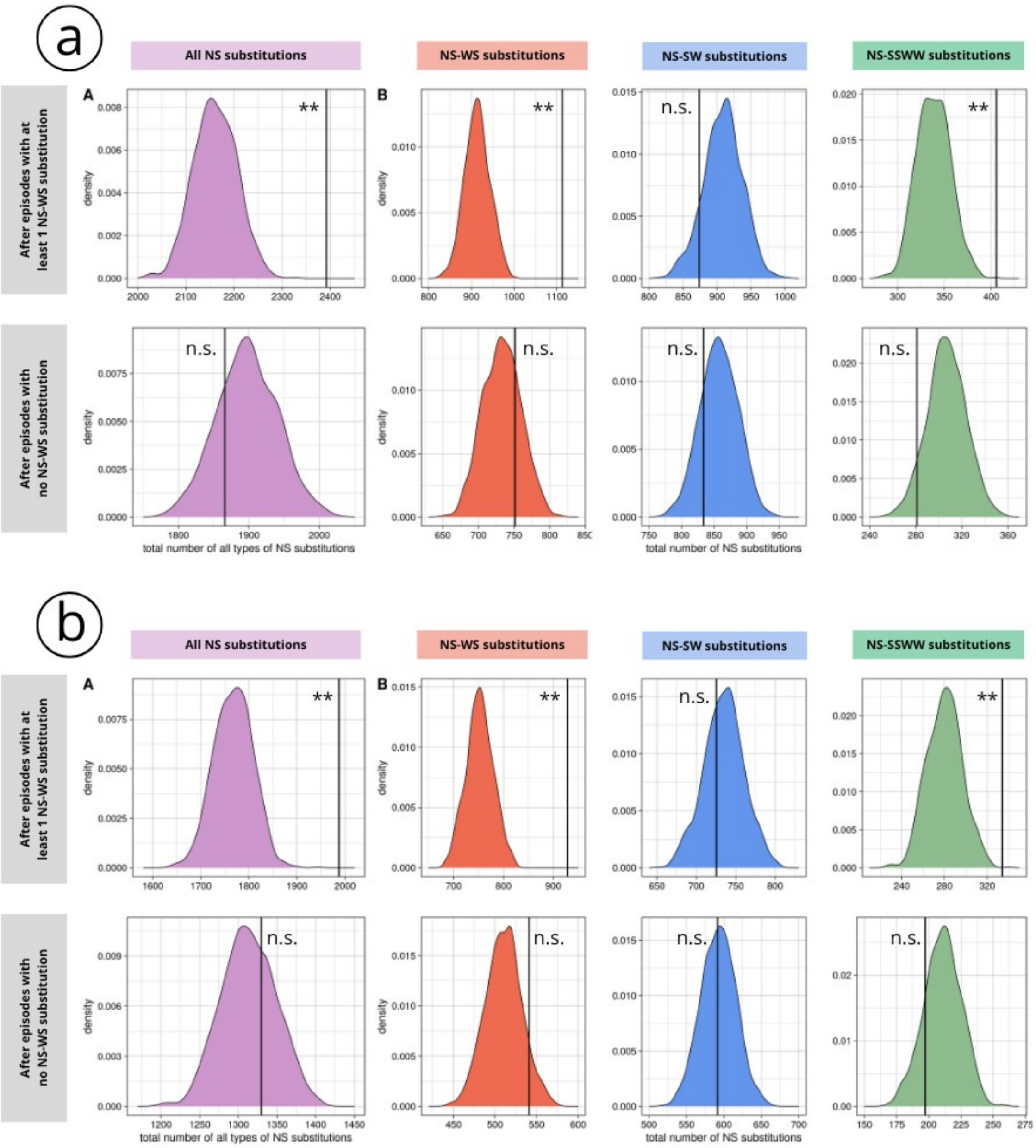


**Figure S4: Compensation signal after "punctual" episodes is still detectable with a more stringent filter on topological discordance**. These figures represent the same as Fig. 5, but with two sets of exons with increasing species tree/exon tree concordance. **a**: filtering threshold at 1.3 (number of episodes with at least one NS-WS=2,087 ; number of episodes with no NS-WS=5,173). **b**: filtering threshold at 1.1 (number of episodes with at least one NS-WS=1,333 ; number of episodes with no NS-WS=2,518).

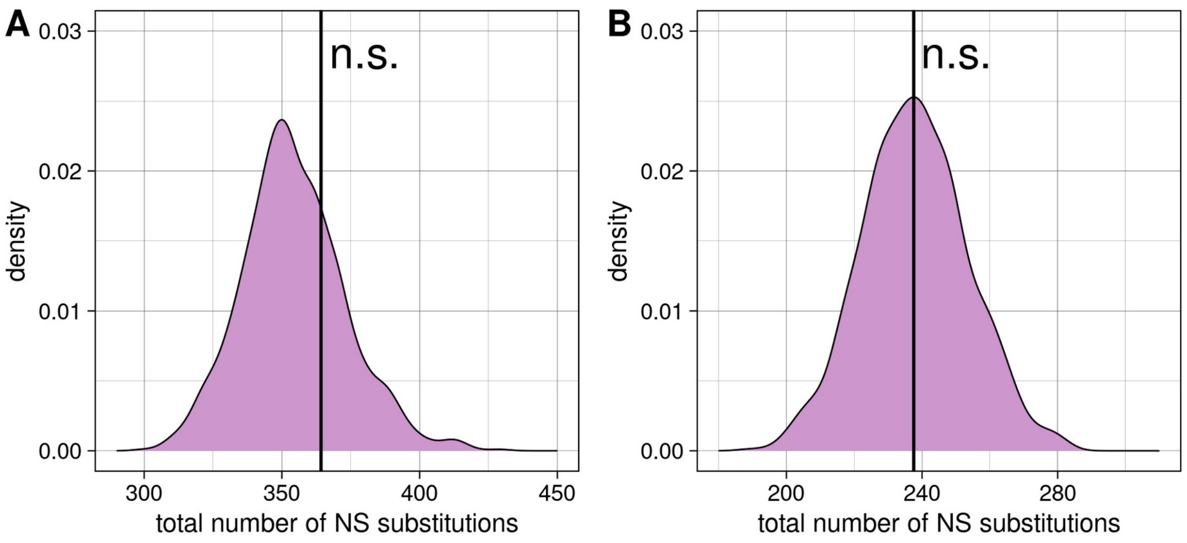


**Figure S5 : No excess of NS substitutions before "punctual" episodes. A :** episodes with at least one NS-WS substitution ; **B :** episodes without any NS-WS substitution. n.s. : p-value >= 0.05.

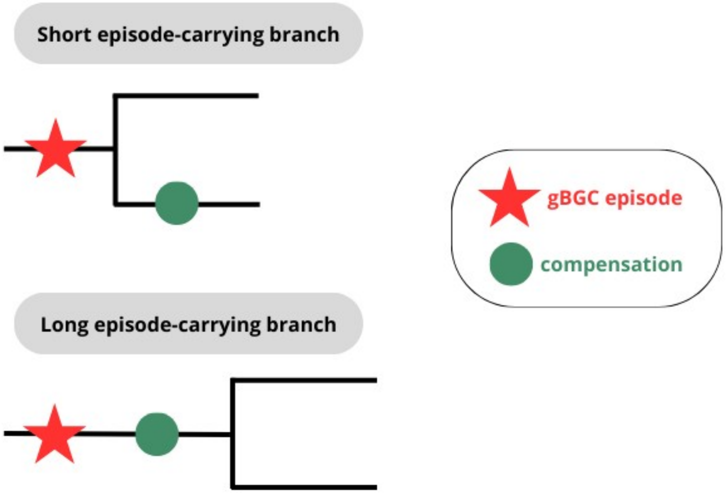


**Figure S6: Compensation probably occurs quickly after gBGC episodes.** The figure illustrates the interpretation of Fig.6, which shows that the compensatory response is detectable only when episode-carrying branches are shorter than 0.006 substitution per site. This suggests that compensation occurs quickly after the episode such that when the episode occurs in a short branch (upper case), compensation occurs and is detectable in the downstream branches; but when the episode occurs in a long branch (lower case), compensation is likely to occur in the same branch, such that no compensation signal is detectable in the downstream branches.

**Supplementary methods: Classifying episodes**

We developed a maximum likelihood (ML) classifier of gBGC episodes in order to distinguish localized, short-lived episodes from large-scale and/or long-lasting effects of gBGC, based on synonymous substitution counts. We model episodes as a mixture of three categories: $C_1$ = punctual, $C_2$ = two-branch, and $C_3$ = all-exons. The punctual category $C_1$ is intended to contain episodes limited to a single exon and a single branch of the tree – our main target in this study. The two-branch category $C_2$ is intended to contain episodes limited to a single exon but also affecting downstream branches. This is expected to happen if a gBGC episode occurs shortly before a speciation event and persists through it. The all-exons category $C_3$ is intended to contain episodes durably affecting the whole gene, as described by Pracana et al. (2020). We call $p_1$, $p_2$ and $p_3$=1-$p_1$-$p_2$ the proportions of punctual, two-branch and all-exons episodes, respectively.

We call "focal exon" the exon of a gene affected by the considered episode, and "other exons" all the other exons of the same gene. We call "focal branch" the branch in which the episode happened, and "descending branches" the two branches immediately downstream the focal branch. We assume that episodes within each of the three categories share a common substitution process, and particularly, a common equilibrium GC-content at neutral sites, θ. In punctual episodes, the synonymous substitution process is described by two parameters, $\theta_1$ and $\theta_1$*, where $\theta_1$* is the equilibrium GC-content for the focal exon in the focal branch, and $\theta_1$ is the equilibrium GC-content for the focal exon in the descending branches, and for other exons, both focal and descending branches. In two-branch episodes, the synonymous substitution process is described by two parameters, $\theta_2$ and $\theta_2$*, where $\theta_2$* is the equilibrium GC-content for the focal exon (focal and descending branches), and $\theta_2$ is the equilibrium GC-content for other exons (focal and descending branches). In all-exons episodes, we have a single parameter, $\theta_3$, which is the equilibrium GC-content for all exons, focal and other, in both the focal and descending branches. The model therefore has seven parameters: the two proportions, $p_1$ and $p_2$, and the five equilibrium GC-content $\theta_1$, $\theta_1$*, $\theta_2$, $\theta_2$*, and $\theta_3$.

For each detected episode *e*, the data consists in the counts of WS and SW synonymous substitutions in the focal and descending branches, for all exons:

$$e = (f_i, d_i) \qquad [1]$$

where $f_i$ = ($f_i$[WS], $f_i$[SW]) stands for the synonymous substitution counts for the $i^{th}$ exon in the focal branch, and $d_i$ = ($d_i$[WS], $d_i$[SW]) in the descending branches. A↔T and C↔G substitutions are here ignored. Below we assume that the considered episode hit the first exon, so that $f_1$ represents the counts for the focal exon, and $f_k$ , $k$>1 the counts for other exons.

The probability of observing a particular set of counts $e$=($f_i$, $d_i$) can be written as a linear combination across the punctual, two-branch and all-exons categories:

$$P(e) = p_1 P_1(e) + p_2 P_2(e) + p_3 P_3(e) \quad [2]$$

where lower-case $p$'s are model parameters standing for the category proportions, and upper case $P$'s are the probability of the data given a category. $P_1(e)$, for instance, is the probability of observing $e$ knowing it belongs to the punctual category. These probabilities are given by the following formulas:

$$P_1(e) = B(f_1,\theta_1^*)\ B(d_1,\theta_1)\ \Pi_{k>1}\ (B(f_k,\theta_1)\ B(d_k,\theta_1)) \quad [3]$$

$$P_2(e) = B(f_1,\theta_2^*)\ B(d_1,\theta_2^*)\ \Pi_{k>1}\ (B(f_k,\theta_2)\ B(d_k,\theta_2)) \quad [4]$$

$$P_3(e) = \Pi_{k>0}\ (B(f_k,\theta_3)\ B(d_k,\theta_3)) \quad [5]$$

where $B(f,\theta)$ is the binomial probability of randomly drawing counts $f$=($f$[WS],$f$[SW]) knowing that equilibrium GC-content is θ. This is given by

$$B(f,\theta) = (f[WS]+f[SW])!\ \gamma^{f[WS]}\ (1-\gamma)^{f[SW]} / (f[WS]!\ f[SW]!) \quad [6]$$

Here γ is the probability that a particular synonymous substitution is of type WS, knowing that equilibrium GC-content is θ. This probability depends on the GC-content at synonymous positions in the considered exon, $g$ :

$$\gamma = \theta(1-g) / (\theta(1-g)+ (1-\theta)g) \quad [7]$$

This results from the fact that, calling $u$ the W→S mutation rate and $v$ the S→W mutation rate, we have $\theta=u/(u+v)$ and $\gamma=u(1-g)/(u(1-g)+vg)$. Rearranging gives equation [7]. $g$ was here approximated as the average GC3 across all available sequences for the considered exon.

The likelihood of the whole data set was obtained by multiplying probabilities $P(e)$ across episodes, assuming that episodes evolve independently. The likelihood was maximized over the parameter space using a modified version of the Newton-Raphson algorithm and ten random starting points. This provided an estimate for each of the seven parameters.

We used a Bayesian empirical approach to classify episodes. Assuming that the ML estimates of parameters correspond to the truth, the posterior probability for episode *e* to belong to category punctual under our model is:

$$P(e \text{ in } C_1) = p_1 P_1(e) / P(e) \quad [8]$$

and similarly for the two-branch and all-exons categories:

$$P(e \text{ in } C_2) = p_2 P_2(e) / P(e) \quad [9]$$

$$P(e \text{ in } C_3) = p_3 P_3(e) / P(e) \quad [10]$$

In addition, we measured for each episode the goodness of fit by comparing $P(e)$ (using the ML parameter estimates) with the probability of the data under a degenerate model that assigns one parameter to each data point. To calculate the degenerate likelihood of each episode, we simply multiplied the binomial probability of equation [6] across all exons and branches assuming that the expected frequency γ equals the observed *f*[WS]/(*f*[WS]+*f*[SW]). Twice the difference in log-likelihood between the degenerate and our model was compared to a $\chi^2$ distribution with $2n_e$ degrees of freedom, where $n_e$ is the number of exons in the considered gene. The resulting *p*-value was used as an episode-specific measure of the goodness of fit. Episodes departing all of our three categories will have a low likelihood under our model, resulting in a large difference in log-likelihood with the degenerate model and a small *p*-value.In this case, we rejected our model and the episode was not considered. Only episodes that were sufficiently well described by the C1 category were kept in the analysis.

A program implementing these calculations was developed in C and is available at *https://github.com/Marie-98/gBGC_comp*
